# Body size-dependent effects on the distribution patterns of phoretic mites of the multi-symbiont *Rhynchophorus ferrugineus* (Olivier, 1790) host

**DOI:** 10.1101/2023.02.06.527267

**Authors:** Inês Matos, Diogo Silva, João Oliveira, Claúdia Gonçalves, Rita Alves, Nuno Pereira, Francisco Catarino, Olga M. C. C. Ameixa, José Américo Sousa, Luis Filipe Rangel, Maria João Santos, Camilo Ayra-Pardo

## Abstract

Phoretic mites have been found attached to different body parts of the red palm weevil (RPW), *Rhynchophorus ferrugineus* (Olivier, 1790), to disperse. However, the question of how the patterns of attachment sites are formed remains intriguing. Here, we conducted the first study of RPW-associated phoretic mites in Portugal, particularly in the districts of Viana do Castelo, Braga, Porto and Aveiro in Northern Portugal (macrohabitat), and investigated the patterns of mite distribution on six body parts of RPW (microhabitat). At the macrohabitat level, we detected seven phoretic mite taxa actively using the RPW host in each of the four studied districts, all documented for the first time in association with this invasive exotic species in Portugal. However, their relative abundance (species evenness) varied between districts, as did species diversity. All examined weevils carried mites, and the prevalence of the different taxa did not differ between districts or sex of weevils. Measured by mean abundance and degree of aggregation, *Centrouropoda* sp. proved to be the common dominant taxon, while *Acarus* sp. And *C. rhynchoporus* were considered common subordinate taxa and *Uroovobella* sp., Mesostigmata, *N. extremica* and *Dendrolaelaps* sp. sparse taxa. At the microhabitat level, all taxa were present in all body parts of the RPW; the highest abundance was in a region encompassing the inner surface of the elytra and the membranous hind wings (subelytral space). Analysis of niche overlap revealed that the distribution patterns of phoretic mite taxa on the RPW were not randomly structured. In the subelytral space, interspecific coexistence of mites increased as a function of body size difference with the dominant *Centrouropoda* sp. We conclude that the distribution patterns of RPW-associated phoretic mites show body size-dependent effects that resulted in the dominant taxon displacing similar size taxa and accepting taxa with which it has the greatest size difference as co-habitants.

## Introduction

The red palm weevil (RPW), *Rhynchophorus ferrugineus* (Olivier, 1790) (Coleoptera: Curculionidae), is a serious pest of palms (Arecaceae) native to South Asia and Melanesia. Since the 1980s, RPW has become an invasive exotic insect in the Euro-Mediterranean region, the Near East, parts of North Africa and the Caribbean (EPPO, 2023). In Portugal, it was first detected in 2007 in the south, in the Algarve region (Fernandes, 2007), and has since spread rapidly throughout the five regions of mainland Portugal and the Madeira Archipelago (Boavida & Franca, 2008; Pete, 2010; Ramos et al., 2013). In the Azores Archipelago, as far as we know, there are no records of its presence. The insect causes economic and ecological impacts after feeding on and destroying the Canary Island date palm *Phoenix canariensis* (Chabaud, 1882), one of its preferred hosts, which is widely used as an ornamental plant in Portuguese coastal cities (Fernandes, 2016). The ability of RPW to grow and develop under a wide range of climatic conditions is mainly attributed to its cryptic feeding behaviour as a borer larva inside the host plant, protected from external temperature and humidity fluctuations (Murphy & Briscoe, 1999; Faleiro, 2006). In addition, the hidden feeding behaviour protects it from natural enemies and pesticides. The recent discovery of RPW damaging sugarcane plants in a field in China shows that RPW can also be highly adaptable to new plant hosts (Qin et al., 2022).

RPW has been described to carry phoretic deutonymph mites (Astigmata, Mesostigmata) on its body (Wisniewski et al., 1992; Longo & Ragusa, 2006; Mesbah et al., 2008; Porcelli et al., 2009; El-Sharabasy, 2010; Mazza et al., 2011; Al-Deeb et al., 2011; Al-Dhafar & Al-Qahtani, 2012; Allam et al., 2013; Dilipkumar et al., 2015; Farahani et al., 2016; Slimane-Kharrat & Ouali, 2019; Abolafia & Ruiz-Cuenca, 2020). This form of temporary symbiotic relationship between mites (phoronts) and the weevil (host) ensures the dispersal of the former (Seeman & Walter, 2023). Some phoretic mite species have specialised, stalk-like temporary structures for attachment, such as the anal pedicel secreted by perianal glands (Bajerlein et al., 2013). Phoretic mites can occur in groups of up to thousands of individuals on specific parts of the host’s body and are thought to impose costs on the host in the form of reduced lifespan (Mazza et al., 2011) or reduced fertility and fecundity of the female (Hodgkin et al., 2010), at least under laboratory conditions. Phoresis has also been considered a precursor to parasitism, as deutonymphs of *Hemisarcoptes cooremani* (Thomas, 1961) have been found to parasitise their dispersal host *Chilocorus cacti* (Linnaeus, 1767) by sucking its body fluids (Houck & Cohen, 1995). Moreover, the phoretic mites *Aegyptus rhynchophorus* Elbishlawi & Allam 2007 and *Aegyptus alhassa* Al-Dhafar & Al-Qahtani 2012 have been found feeding on different developmental stages of RPW and have been proposed for pest control under field conditions (Allam & El-Bishlawi, 2010; Al-Dhafar & Al-Qahtani, 2012; Allam et al., 2013; Allam & Elbadawy, 2017). Other phoretic mites are nematophagous (phagophillia) (Cakmak et al., 2013; Stirling et al., 2017), which may reduce the efficacy of entomopathogenic nematodes used to treat RPW-infested palms in Portugal and other countries (DGAV, 2013). The study of RPW-associated phoretic mites can therefore help to identify the biodiversity components of the ecosystem in which this invasive exotic weevil has become established and to develop potential biological control measures against this palm pest.

Most studies on RPW-associated phoretic mites have documented that coexisting mite species form distribution patterns on RPW body regions; however, how these patterns emerge is intriguing and not yet fully understood. It has been proposed that interspecific overlap in space utilisation (niche overlap) is the cause of interspecific interactions that shape species distribution patterns and lead to species coexistence (Holt, 2009). Niche overlap analyses have been used to infer the extent of such interactions by statistically comparing observed values with expected values generated by randomisation with null-models representing assemblages in which no biological mechanisms regulate species coexistence (Gotelli & Graves, 1996).

Rohde (1984) has defined the habitat for parasites on two levels: the microhabitat and the macrohabitat, and this can also apply to phoretic organisms. According to this author, the macrohabitat is the environment in which the host lives and which is controlled by physical and chemical parameters. The microhabitat is the host itself, i.e. the confined environment in which the parasite lives and settles. Several studies have been conducted on parasites at these two levels (Castro & Santos, 2013; Gobbin et al., 2021). In the RPW-phoretic mite association, the macrohabitat of the mite includes the environment where the RPW, the host, lives (as a geographical distribution), and its microhabitat consists of the preferred regions on the host’s body. To our knowledge, no studies combining macrohabitat and microhabitat analyses of RPW-associated phoretic mites have been conducted. In the present study, we documented for the first time the diversity of RPW-associated phoretic mite species in Portugal, particularly in the districts of Viana de Castelo, Braga, Porto and Aveiro in Northern Portugal (macrohabitat), and their distribution in different parts of the weevil’s body (microhabitat). The generated dataset of mite abundance per body part was then analysed for niche overlap using null models to reveal possible mechanisms of coexistence of mixed mite taxa in this multi-symbiont host.

## Materials and Methods

### Insect collection and sample preparation

From July 2021 to March 2022, Picusan traps (Koppert, Berkel en Rodenrijs, The Netherlands) containing RPW-specific synthetic aggregation pheromones (Ao Midori Biocontrol S.L., Barcelona, Spain) were placed at eight different sites in the Northern Portugal districts of Viana do Castelo, Braga, Porto and Aveiro near visibly infected Canary Island date palms. The traps were placed at ground level to attract more weevils (Hallett et al., 1999); neighbouring traps were placed 25 metres apart. The name and geographical localisation of this eight sites are: Moledo do Minho (41.847698, -8.864375) in Viana do Castelo; Famalicão (41.361287, - 8.539348) in Braga; Marquês (41.161283, -8.604427), Porto Botanical Garden (41.153008, - 8.642730) and Foz do Douro (41.147716, -8.670845) in Porto; Esmoriz (40.955908, -8.650886), Santiago Campus - University of Aveiro (40.635053, -8.659500) and Barra Beach (40.632630, - 8.748307) in Aveiro. The traps were inspected weekly; the captured weevils were transported to the laboratory in plastic containers and fed with apple slices. The specimens were then stored in the refrigerator at 8 °C until dissection.

### Identification and distribution patterns of RPW-associated mites

In the laboratory, weevils were separated by sex and dissected individually under a stereomicroscope. Phoretic mites were counted on six body parts of the weevils, i.e. neck, head-antenna, thorax, legs, ventral surface (abdominal sterna), and subelytral space (membranous hind wings + inner elytra surface), then removed with a camel hair brush and preserved in 70% ethanol.

For identification, selected specimens of each observed mite morph were mounted on microscopic slides in lactic acid (90% solution in water) (Helle & Sabelis, 1985) and examined with an Axiophot microscope (Carl Zeiss, Oberkochen, Germany) connected to a computer running Leica Application Suite X (LAS X) image processing software. Mites were identified to the lowest possible taxonomic level based on the original descriptions and illustrations in which these taxa were first described (Womersley, 1954; McGraw & Farrier, 1969; Griffiths, 1970; Fain, 1974; Lindquist, 1975; Kinn, 1984; Wisniewski et al., 1992; Krantz & Walter, 2009; Porcelli et al., 2009; Farahani et al., 2016; Abo-Shnaf & Allam, 2019) and the length of their ventral idiosoma measured.

### Statistical methods

All analyses were conducted using the open-source R environment (R developmental core team, 2021). Prevalence (the proportion of weevils that harboured mites), mean intensity (the mean number of mites per infected weevil), mean abundance (the mean number of mites per weevil) and their respective confidence intervals (at a 95% confidence level) were calculated as in Rózsa et al. (2000). The prevalence of mite species in each district and between female and male weevils was compared using Fisher’s exact test, with the exact *P* value given. Kruskal Wallis chi-squared in conjunction with Dunn’s *post-hoc* test (Bonferroni correction applied) at *P* < 0.05 as significance level were performed to detect differences between districts in the mean intensity of each mite species (Rózsa et al. 2000). Poulin’s index of discrepancy (D) (Poulin, 1993) - a measure of intraspecific aggregation - was calculated as in Morrill et al. (2022), using the bootstrap method to obtain 95% confidence intervals based on 2000 replicates. Alpha diversity calculations in each mite assemblage (Viana do Castelo, Braga, Porto and Aveiro) included abundance-based Hill diversity numbers (^*q*^D) of the three *q* orders, i.e. *q*=0 (species richness), *q*=1 (Shannon diversity or effective number of frequent species in the assemblage) and *q*=2 (Simpson diversity or effective number of highly frequent species in the assemblage) and were performed using the R package iNEXT (Chao et al., 2014; Hsieh et al., 2016; Chao et al., 2020). Plotting of the Lorenz curve and calculation of the Gini index were performed with the R package ineq (Zeileis & Kleiber, 2014). While the Lorenz curve is a graphical representation of the inequality between the size of individuals in a community (species evenness), the Gini index measures the extent to which the size of individuals within a community deviates from a perfectly equal distribution and ranges from 0 (complete equality) to 1 (complete inequality) (Patil & Taillie, 1979). To visualise the distribution of mites on the host and to evaluate the uncertainty of the hierarchical cluster analysis (method “ward.D”, “Euclidean” distance and bootstrap replications set to 10000), a resource utilisation matrix consisting of the number of individual occurrences of each taxa (species abundance) in the different body parts of RPW (microhabitats) was analysed using the R-packages bipartite (Dormann, 2022) and pvclust (Suzuki & Shimodaira, 2006), respectively. The analysis of niche overlap was performed with the R package EcoSimR 1.00 (Gotelli et al., 2015). First, the aforementioned 7×6 resource utilisation matrix was used to determine the Pianka’s index of niche overlap, which represents the average mean spatial overlap of all possible species pairs in the community and ranges from 0 (no overlap) to 1 (complete overlap) (Pianka, 1974). Subsequently, the observed Pianka’s index was statistically compared (*P* < 0.05) with expected values from simulated 7×6 resource utilisation matrices generated using the Monte Carlo randomisation algorithms RA2 and RA3 for null model analysis (Gotelli & Graves, 1996). RA2 assumes random equal utilisation of space (i.e. relaxed niche breadth), while RA3 assumes that each body part has an equal probability of harbouring a species (i.e. retaining niche breadth) (Lawlor, 1980). The Pearson coefficient (r) with *P* < 0.05 was used for all correlation analyses.

## Results

### Macrohabitat level — Diversity and ecology of RPW-associated phoretic mites in Northen Portugal

A total of 238 adult RPW were collected with pheromone traps in four districts of the Northern Portugal region. The sample was distributed as follows: 30 in the district of Viana do Castelo, 53 in Braga, 44 in Porto and 111 in Aveiro with female:male ratios of 1.5:1, 1.4:1, 2.1:1 and 2.2:1, respectively. In this sample, 100% of the captured weevils carried phoretic deutonymphs of seven mite taxa documented for the first time as associated with the invasive exotic RPW in Portugal; these were the species *Nenteria extremica* Kontschan, Mazza, Nannelli & Roversi 2014 (Mesostigmata: Trematuridae) and *Curculanoetus rhynchophorus* Fain 1974 (Astigmata: Histiostomatidae), and five unspecified taxa, namely *Centrouropoda* sp. Berlese 1916 (Mesostigmata: Uropodidae), *Uroobovella* sp. Berlese 1905 (Mesostigmata: Urodinychidae), *Acarus* sp. Linnaeus 1758 (Acariformes: Acaridae), *Dendrolaelaps* sp. Halbert 1915 (Mesostigmata: Digamasellidae), and one from the order Mesostigmata. The order of size of the identified mites by length of idiosoma ± standard error was *Uroobovella* sp. (886.3 ± 30.9 μm) > *Centrouropoda* sp. (614.2 ± 29.2 μm) > *N. extremica* (394.8 ± 16.4 μm) > Mesostigmata (340.3 ± 8.3 μm) > *Dendrolaelaps* sp. (296 ± 15.7 μm) > *C. rhynchophorus* (239.1 ± 16.3 μm) > *Acarus* sp. (199.7 ± 16.1 μm) (**Fig. 1**).

**Figure 1.**
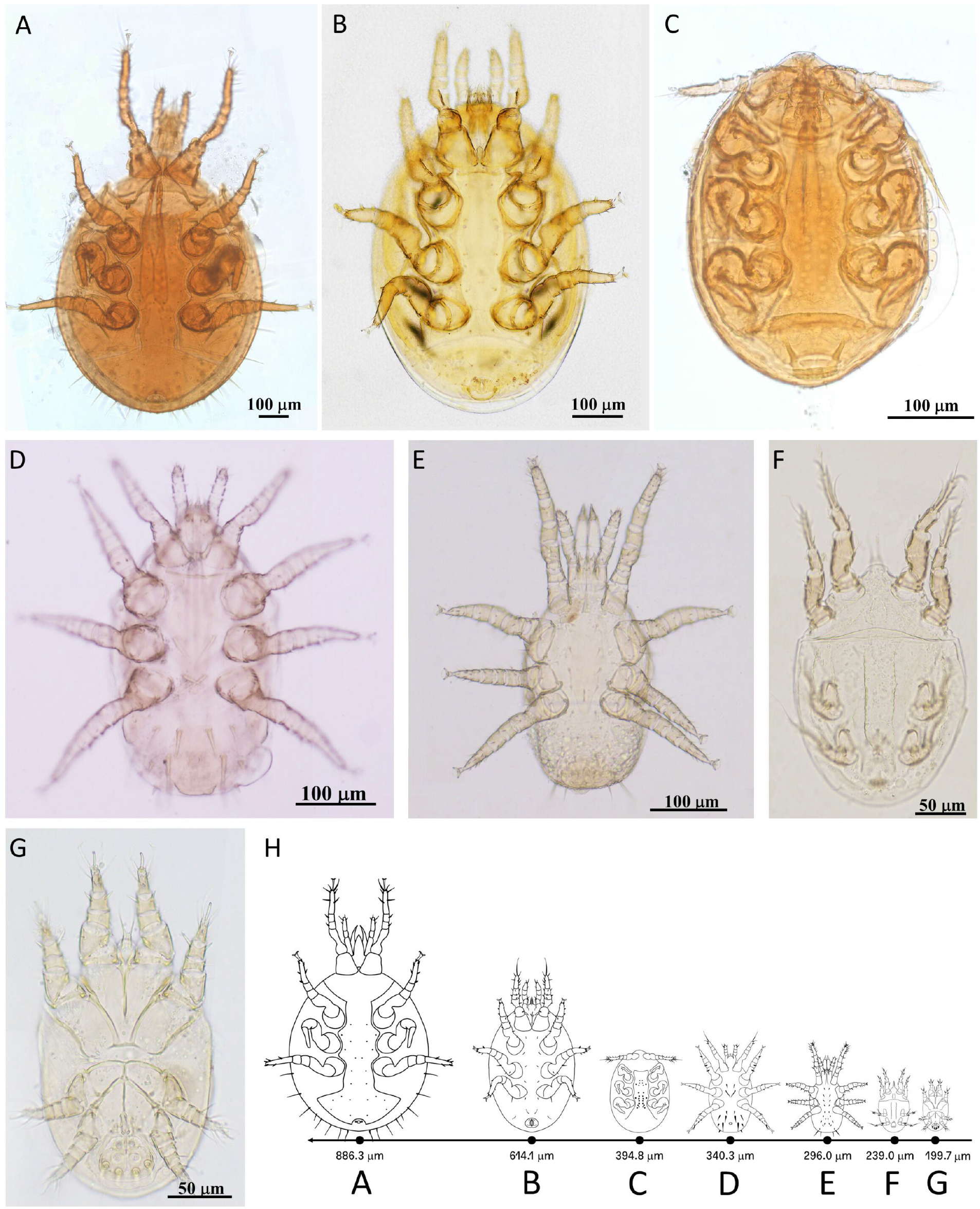
Ventral views of phoretic deutonymphs associated with the red palm weevil in Northern Portugal’s districts of Viana do Castelo, Braga, Porto and Aveiro. The mites were identified using an Axiophot microscope (Carl Zeiss, Oberkochen, Germany) connected to a computer running Leica Application Suite X (LAS X) image processing software. The length of the ventral idiosoma ± standard deviation was determined for each mite species (*n*=10). (A) *Uroobovella* sp. (886.3 ± 30.9 μm), (B) *Centrouropoda* sp. (614.2 ± 29.2 μm), (C) *Nenteria extremica* (394.8 ± 16.4 μm), (D) Mesostigmata (340.3 ± 8.3 μm) (E) *Dendrolaelaps* sp. (296 ± 15.7 μm), (F) *Curculanoetus rhynchophorus* (239.1 ± 16.3 μm), (G) *Acarus* sp. (199.7 ± 16.1 μm). (H) Schematic drawing with the identified deutonymphs at the same scale.

Species diversity in the four RPW-associated mite assemblages (i.e. Viana do Castelo, Braga, Porto, Aveiro) was compared using rarefaction/extrapolation curves of Hill numbers ^*q*^*D* (Chao et al., 2014; Chao et al., 2020), with exponential value *q*=0 for richness, *q*=1 for Shannon’s diversity and *q*=2 for Simpson’s diversity (**Fig. 2A**). We found a steep increase up to seven accumulated taxa in all districts; however, Aveiro needed more individuals to reach the seven taxa compared to Viana do Castelo, Braga and Porto. Shannon’s diversity, which gives more weight to common taxa in the assemblage, was well recorded in all districts, being significantly higher in Viana do Castelo and decreasing in Porto, Aveiro and Braga with non-overlapping confidence intervals. Simpson’s diversity, which counts the effective number of highly common taxa in the assemblage, was also well recorded in all districts; Viana do Castelo and Porto had similarly high values with overlapping 95% confidence intervals on the rarefaction/extrapolation curves, while Aveiro had a lower value and Braga the lowest value. The four mite assemblages were then compared in terms of species evenness — a measure of the relative abundance of species within the community — using the Lorenz curve method and calculating the Gini index (**Fig. 2B**). In combination with species richness, these tools can be used to assess species diversity. Accordingly, species evenness decreased in the order Viana do Castelo>Porto>Aveiro>Braga (**Fig. 2B**), confirming that the assemblage Viana do Castelo had the most even distribution and thus the greatest diversity of RPW-associated mite taxa.

**Figure 2.**
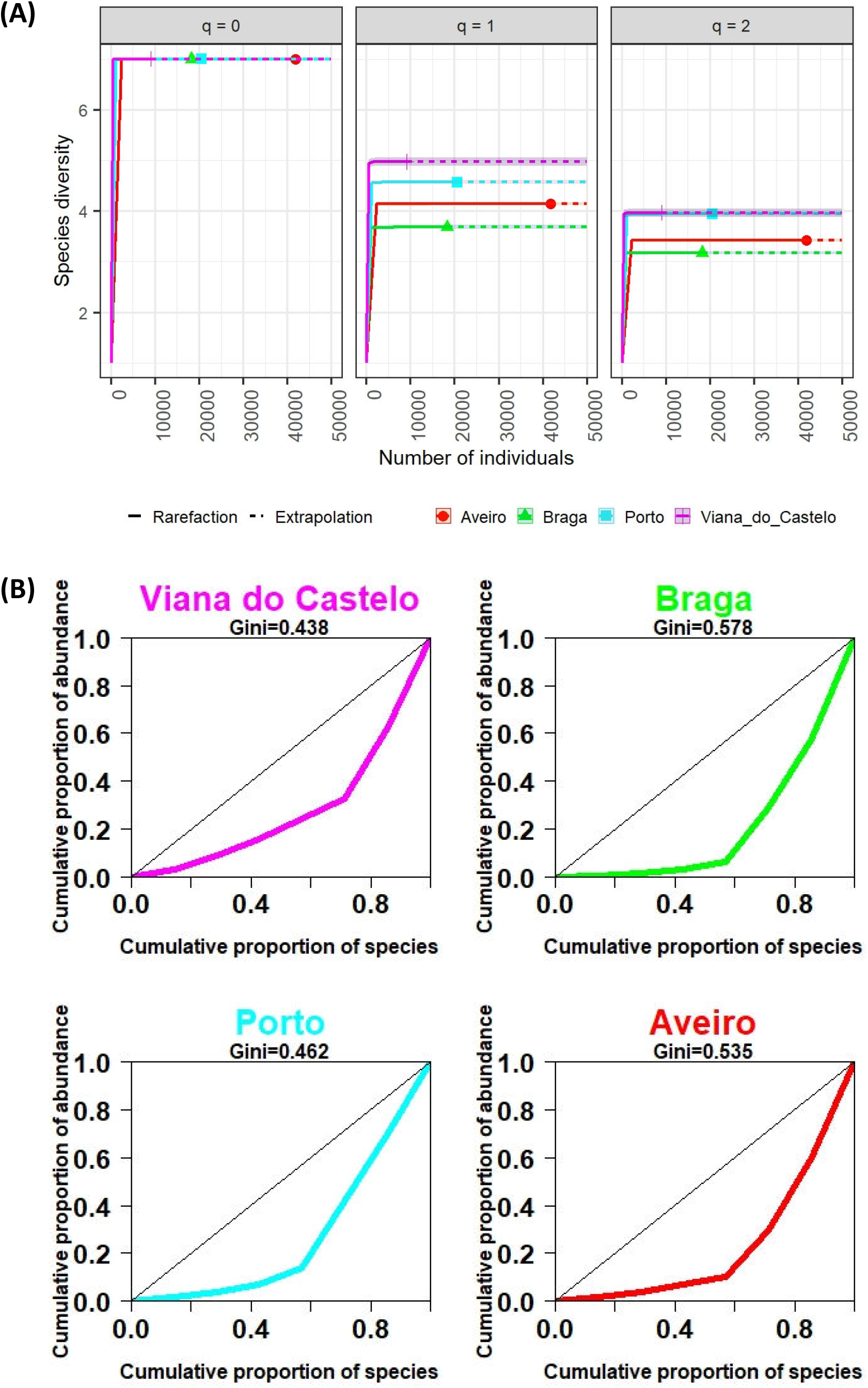
(A) Sample-size-based rarefaction and extrapolation diversity curves for red palm weevil-associated phoretic mite assemblages in Northern Portugal’s districts of Viana do Castelo (magenta), Braga (green), Porto (cyan) and Aveiro (red), based on Hill’s order (*q*) with *q* = 0 (species richness), *q* = 1 (exponential of Shannon entropy index) and *q* = 2 (inverse of Simpson index) (Chao et al., 2020). The solid line is the rarefaction curve and the dotted line is the extrapolation curve. The intersection between the solid and dotted lines represents the observed values. The extrapolation goes up to 50000 individuals to exceed the actual sample size. The shaded areas represent 95% confidence intervals obtained by the bootstrap method based on 100 replicates. (B) Lorenz curves and Gini index comparing the evenness of the relative abundance of red palm weevil-associated phoretic mite assemblages in the Northern Portugal’s districts of Viana do Castelo, Braga, Porto and Aveiro.

The ecology of the four assemblages — in terms of descriptive statistics, such as prevalence, mean intensity, mean abundance and Poulin’s index of discrepancy for each RPW-associated mite taxa — is summarised in **Table 1**. The prevalence (the proportion of weevils that contained mites) of each mite taxa showed no significant differences between districts; it ranged from 88.7% to 83.3% for *Centrouropoda* sp. (*P=*0.877), 66.7% to 44.1% for *N. extremica* (*P=*0.067), 72.7% to 56.6 for *Acarus* sp. (*P=*0.214), 37.7% to 25% for Mesostigmata (*P=*0.561), 50% to 32.4% for *Dendrolaelaps* sp. (*P=*0.277), 30.6% to 15.9% for *Uroobovella* sp. (*P=*0.234), and 48.6% to 36.7% for *C. rhynchophorus* (*P=*0.644). The prevalence of each mite taxa also showed no differences between male and female weevils (*Centrouropoda* sp., *P*=0.88; *N. extremica, P*=0.35; *Acarus* sp., *P*=0.68; Mesostigmata, *P*=0.25; *Dendrolaelaps* sp., *P*=0.67; *Uroobovella* sp., *P*=0.41; *C. rhynchophorus, P*=0.78). The mean intensity (the mean number of mites per infected weevil) of each mite taxa differed significantly between districts only for Mesostigmata (H=12.92, degrees of freedom=3, *P*=0.005; **Table 1**) and *C. rhynchophorus* (H=9.22, degrees of freedom=3, *P* =0.027; **Table 1**). Pearson correlation analysis of the linear relationship between mean abundance (MA; the mean number of mites per weevil) and the Poulin’s index of discrepancy (D; a measure of intraspecific aggregation) of the seven mite taxa in the four mite assemblages showed a highly significant, strong negative correlation [r(28)= - 0.81, *P*<0.0001] (**Fig. 3**). *Centrouropoda* sp. was clearly the dominant taxon with the highest MA values and the lowest D values, followed by *Acarus* sp. and *C. rhynchophorus* (the smallest mites of the seven taxa) with medium to high MA and D values, which were considered common subordinate taxa. The remaining four taxa, i.e. *Uroobovella* sp., *N. extremica*, Mesostigmata, and *Dendrolaelaps* sp., were each found in very low numbers and highly aggregated in the four RPW-associated mite assemblages and were then considered sparse taxa.

**Table 1.**
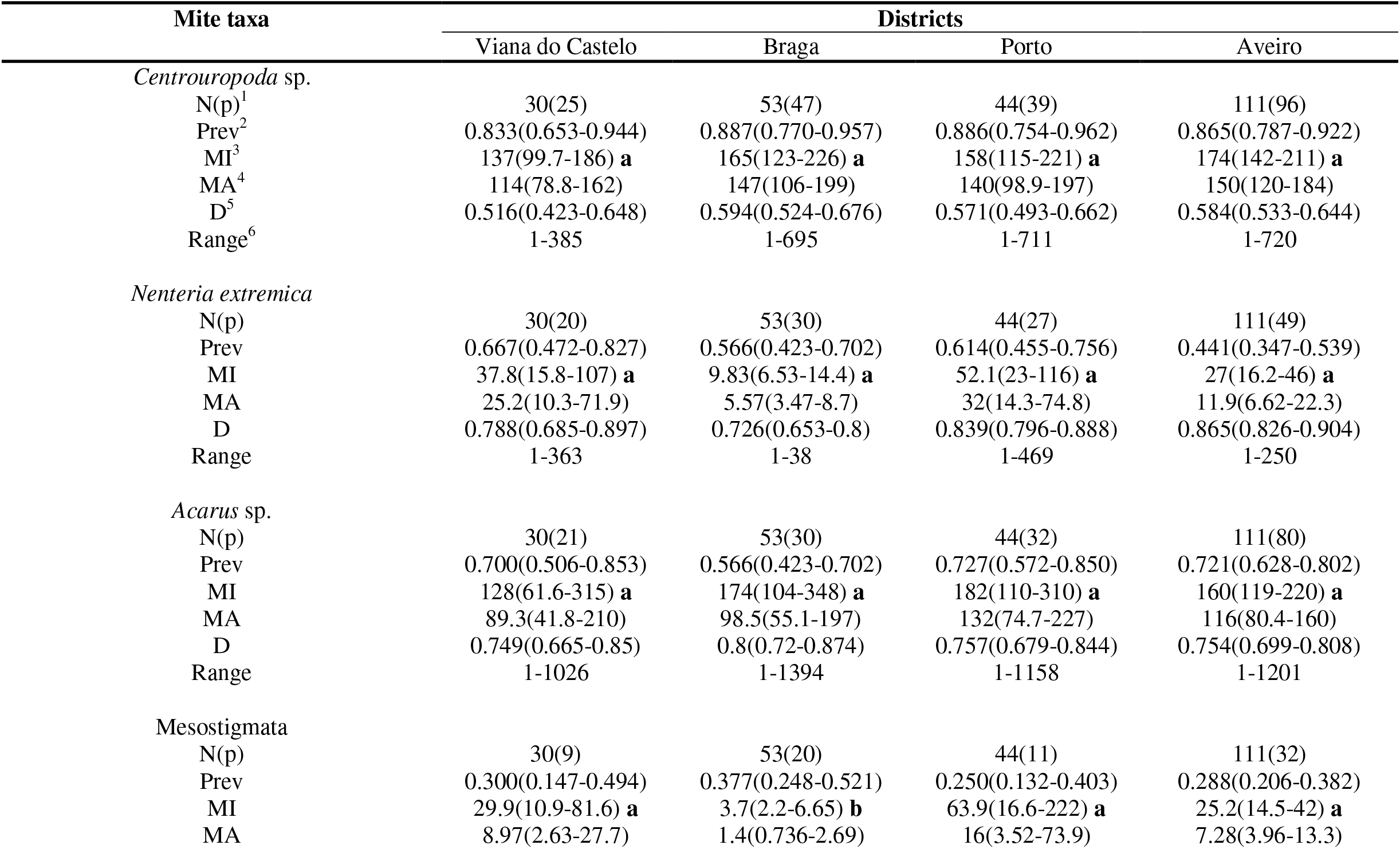
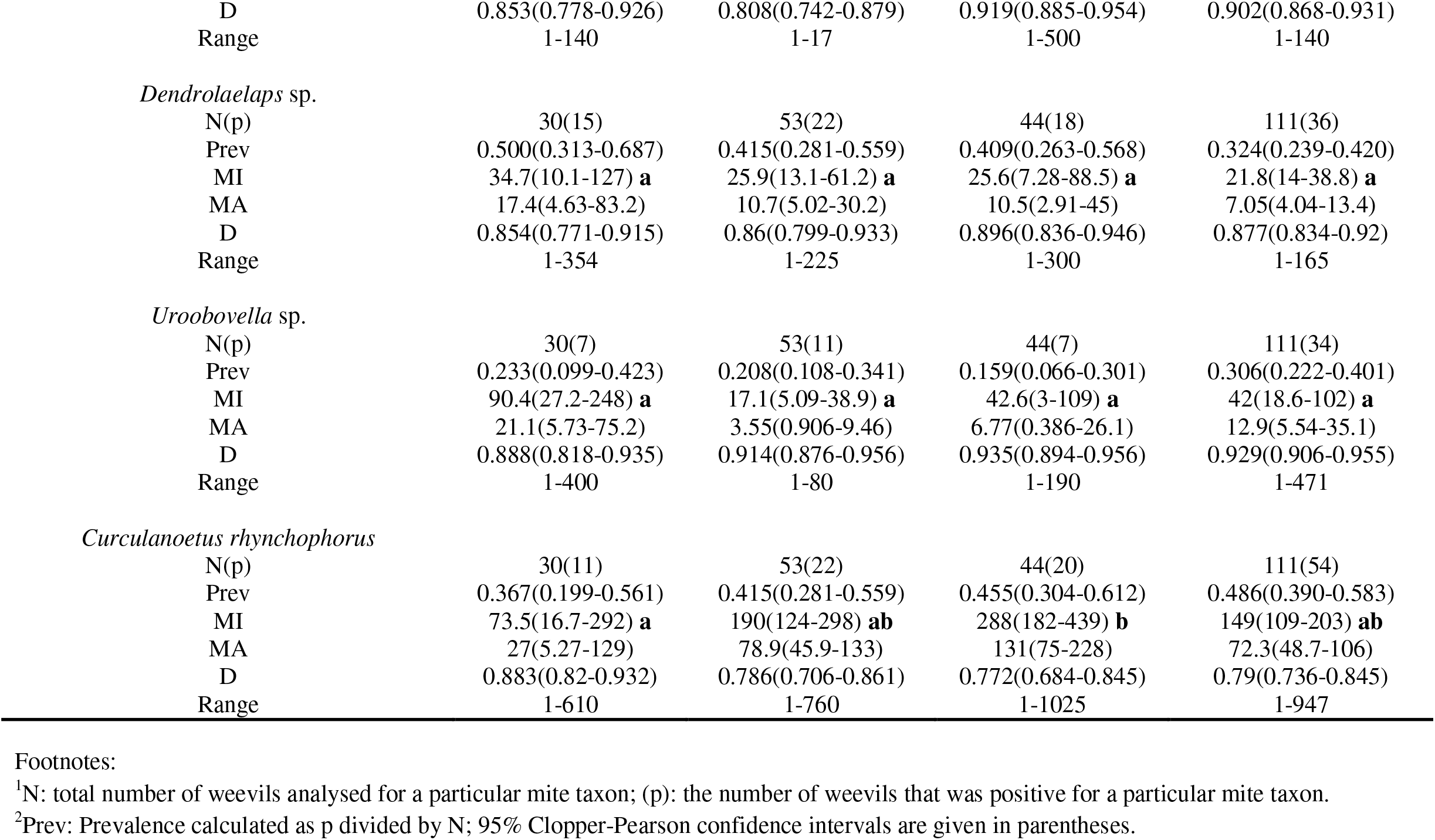
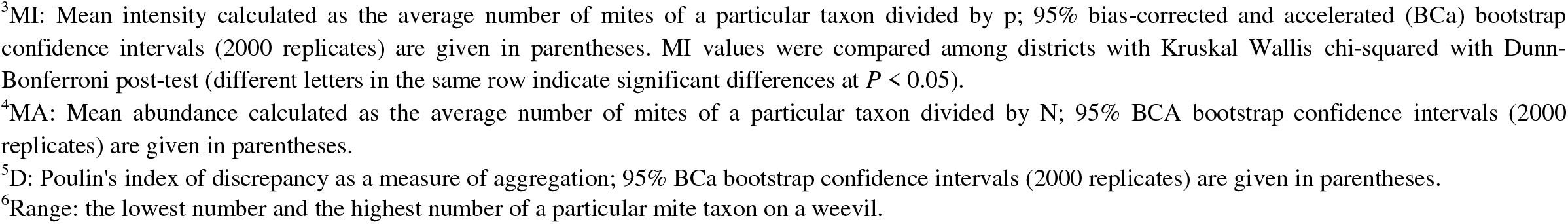
Summary of descriptive statistics of RPW-associated phoretic mites in four districts of Northern Portugal.

| Mite taxa | Districts |  |  |  |
| --- | --- | --- | --- | --- |
|  | Viana do Castelo | Braga | Porto | Aveiro |
| *Centrouropoda* sp. |  |  |  |  |
| N(p)<sup>1</sup> | 30(25) | 53(47) | 44(39) | 111(96) |
| Prev<sup>2</sup> | 0.833(0.653-0.944) | 0.887(0.770-0.957) | 0.886(0.754-0.962) | 0.865(0.787-0.922) |
| MI<sup>3</sup> | 137(99.7-186) **a** | 165(123-226) **a** | 158(115-221) **a** | 174(142-211) **a** |
| MA<sup>4</sup> | 114(78.8-162) | 147(106-199) | 140(98.9-197) | 150(120-184) |
| D<sup>5</sup> | 0.516(0.423-0.648) | 0.594(0.524-0.676) | 0.571(0.493-0.662) | 0.584(0.533-0.644) |
| Range<sup>6</sup> | 1-385 | 1-695 | 1-711 | 1-720 |
| *Nenteria extremica* |  |  |  |  |
| N(p) | 30(20) | 53(30) | 44(27) | 111(49) |
| Prev | 0.667(0.472-0.827) | 0.566(0.423-0.702) | 0.614(0.455-0.756) | 0.441(0.347-0.539) |
| MI | 37.8(15.8-107) **a** | 9.83(6.53-14.4) **a** | 52.1(23-116) **a** | 27(16.2-46) **a** |
| MA | 25.2(10.3-71.9) | 5.57(3.47-8.7) | 32(14.3-74.8) | 11.9(6.62-22.3) |
| D | 0.788(0.685-0.897) | 0.726(0.653-0.8) | 0.839(0.796-0.888) | 0.865(0.826-0.904) |
| Range | 1-363 | 1-38 | 1-469 | 1-250 |
| *Acarus* sp. |  |  |  |  |
| N(p) | 30(21) | 53(30) | 44(32) | 111(80) |
| Prev | 0.700(0.506-0.853) | 0.566(0.423-0.702) | 0.727(0.572-0.850) | 0.721(0.628-0.802) |
| MI | 128(61.6-315) **a** | 174(104-348) **a** | 182(110-310) **a** | 160(119-220) **a** |
| MA | 89.3(41.8-210) | 98.5(55.1-197) | 132(74.7-227) | 116(80.4-160) |
| D | 0.749(0.665-0.85) | 0.8(0.72-0.874) | 0.757(0.679-0.844) | 0.754(0.699-0.808) |
| Range | 1-1026 | 1-1394 | 1-1158 | 1-1201 |
| Mesostigmata |  |  |  |  |
| N(p) | 30(9) | 53(20) | 44(11) | 111(32) |
| Prev | 0.300(0.147-0.494) | 0.377(0.248-0.521) | 0.250(0.132-0.403) | 0.288(0.206-0.382) |
| MI | 29.9(10.9-81.6) **a** | 3.7(2.2-6.65) **b** | 63.9(16.6-222) **a** | 25.2(14.5-42) **a** |
| MA | 8.97(2.63-27.7) | 1.4(0.736-2.69) | 16(3.52-73.9) | 7.28(3.96-13.3) |
| D | 0.853(0.778-0.926) | 0.808(0.742-0.879) | 0.919(0.885-0.954) | 0.902(0.868-0.931) |
| Range | 1-140 | 1-17 | 1-500 | 1-140 |
| *Dendrolaelaps* sp. |  |  |  |  |
| N(p) | 30(15) | 53(22) | 44(18) | 111(36) |
| Prev | 0.500(0.313-0.687) | 0.415(0.281-0.559) | 0.409(0.263-0.568) | 0.324(0.239-0.420) |
| MI | 34.7(10.1-127) **a** | 25.9(13.1-61.2) **a** | 25.6(7.28-88.5) **a** | 21.8(14-38.8) **a** |
| MA | 17.4(4.63-83.2) | 10.7(5.02-30.2) | 10.5(2.91-45) | 7.05(4.04-13.4) |
| D | 0.854(0.771-0.915) | 0.86(0.799-0.933) | 0.896(0.836-0.946) | 0.877(0.834-0.92) |
| Range | 1-354 | 1-225 | 1-300 | 1-165 |
| *Uroobovella* sp. |  |  |  |  |
| N(p) | 30(7) | 53(11) | 44(7) | 111(34) |
| Prev | 0.233(0.099-0.423) | 0.208(0.108-0.341) | 0.159(0.066-0.301) | 0.306(0.222-0.401) |
| MI | 90.4(27.2-248) **a** | 17.1(5.09-38.9) **a** | 42.6(3-109) **a** | 42(18.6-102) **a** |
| MA | 21.1(5.73-75.2) | 3.55(0.906-9.46) | 6.77(0.386-26.1) | 12.9(5.54-35.1) |
| D | 0.888(0.818-0.935) | 0.914(0.876-0.956) | 0.935(0.894-0.956) | 0.929(0.906-0.955) |
| Range | 1-400 | 1-80 | 1-190 | 1-471 |
| *Curculanoetus rhynchophorus* |  |  |  |  |
| N(p) | 30(11) | 53(22) | 44(20) | 111(54) |
| Prev | 0.367(0.199-0.561) | 0.415(0.281-0.559) | 0.455(0.304-0.612) | 0.486(0.390-0.583) |
| MI | 73.5(16.7-292) **a** | 190(124-298) **ab** | 288(182-439) **b** | 149(109-203) **ab** |
| MA | 27(5.27-129) | 78.9(45.9-133) | 131(75-228) | 72.3(48.7-106) |
| D | 0.883(0.82-0.932) | 0.786(0.706-0.861) | 0.772(0.684-0.845) | 0.79(0.736-0.845) |
| Range | 1-610 | 1-760 | 1-1025 | 1-947 |

<sup>1</sup>N: total number of weevils analysed for a particular mite taxon; (p): the number of weevils that was positive for a particular mite taxon.
<sup>2</sup>Prev: Prevalence calculated as p divided by N; 95% Clopper-Pearson confidence intervals are given in parentheses.
<sup>3</sup>MI: Mean intensity calculated as the average number of mites of a particular taxon divided by p; 95% bias-corrected and accelerated (BCa) bootstrap confidence intervals (2000 replicates) are given in parentheses. MI values were compared among districts with Kruskal Wallis chi-squared with Dunn- Bonferroni post-test (different letters in the same row indicate significant differences at $P < 0.05$ ). <sup>4</sup>MA: Mean abundance calculated as the average number of mites of a particular taxon divided by N; 95% BCA bootstrap confidence intervals (2000 replicates) are given in parentheses. <sup>5</sup>D: Poulin's index of discrepancy as a measure of aggregation; 95% BCa bootstrap confidence intervals (2000 replicates) are given in parentheses. <sup>6</sup>Range: the lowest number and the highest number of a particular mite taxon on a weevil.

**Figure 3.**
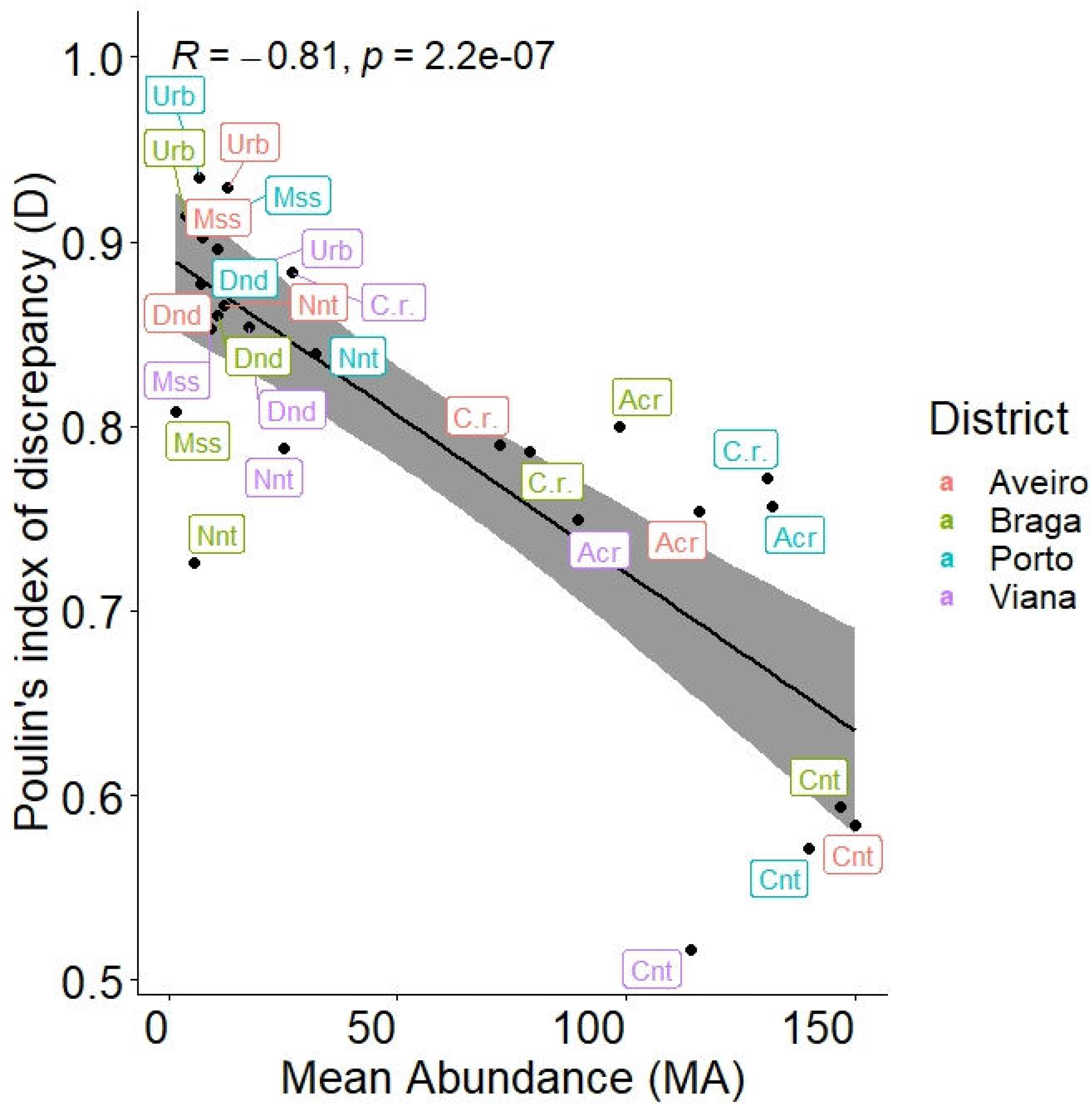
Correlation between Poulin’s index of discrepancy (D) and mean abundance of red palm weevil-associated phoretic mites. The shaded area along the regression line corresponds to the 95% confidence intervals. The inset shows the values of Person’s correlation coefficient (r) and *P*. Significance *P* <0.05. Cnt: *Centrouropoda* sp.; Nnt: *Nenteria extremica*; Acr: *Acarus* sp.; Mss: Mesostigmata; Dnd: *Dendrolaelaps* sp.; Urb: *Uroobovella* sp.; C.r.: *Curculanoetus rhynchophorus*.

### Microhabitat level — Distribution patterns of RPW-associated mites and spatial overlap between co-occurring taxa

Mites were directly separated from six body parts of RPW during identification, namely the head-antenna, neck, legs, thorax, abdominal sterna and subelytral space (membranous hind wings + inner elytra surface) and quantified (**Fig. 4A**). A two-sided plot of mite distribution patterns on the RPW showed that all taxa interacted with the six body parts examined; however, the subelytral space had the greatest number of interactions, which were most pronounced for *Centrouropoda* sp., *C. rhynchophorus* and *Acarus* sp., in that order (**Fig. 4B**). A hierarchical cluster analysis of the distribution patterns of RPW-associated phoretic deutonymphs divided the seven taxa into two main branches, with the common dominant taxon *Centrouropoda* sp. and the common subordinate taxa *C. rhynchophorus* and *Acarus* sp. in the same group and the sparse taxa in a separate group (**Fig. 4C**).

**Figure 4.**
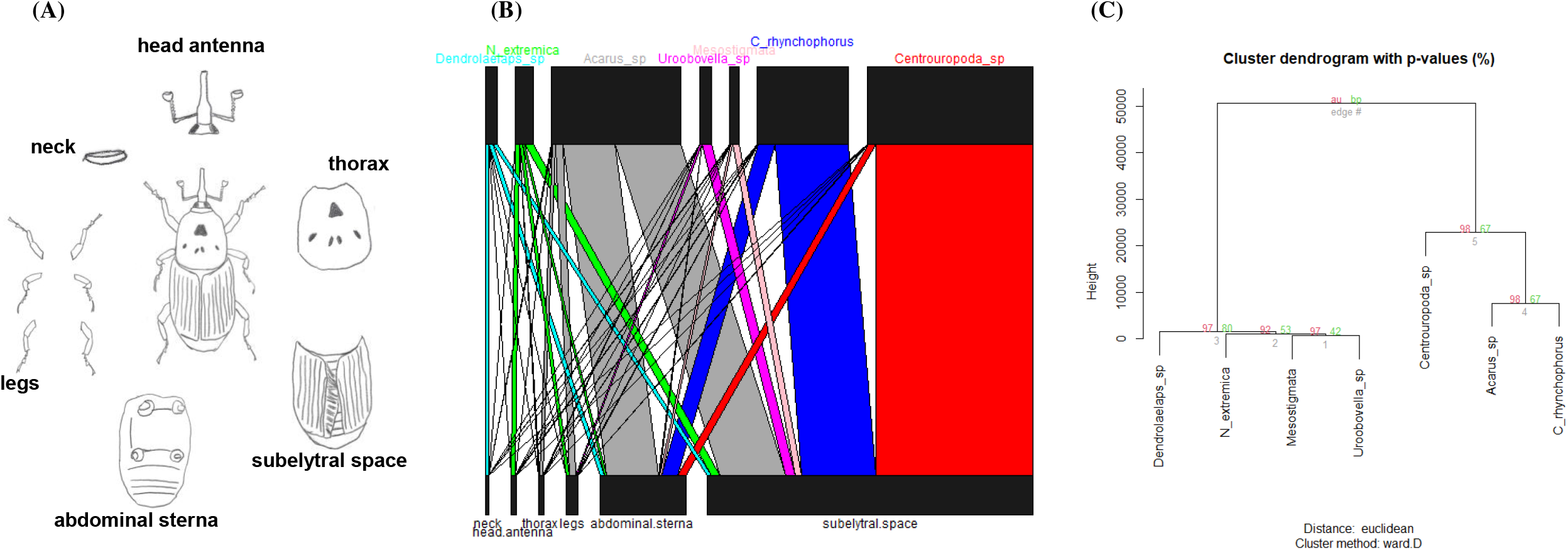
(A) Schematic drawing of the six body parts of the red palm weevil examined for the presence of phoretic mites, i.e. neck, head-antenna, thorax, legs, abdominal sterna, subelytral space (membranous hind wings + inner elytra surface). (B) Bipartite graph illustrating the interaction between phoretic mite taxa (top) and body parts of the red palm weevil (bottom) (7 mite taxa × 6 body parts matrix). Interactions between top and bottom are represented by edges coloured according to the mite taxon involved, with thickness proportional to the number of interactions. (C) Dendogram showing hierarchical clustering of the 7 mite taxa × 6 body parts matrix, performed with the R package pvclust. Values on the branches are approximate unbiased (AU) *P*-values (red, left) and bootstrap probability (BP) (green, right), expressed as percentages (Suzuki and Shimodaira, 2006). Clustering method: ward.D (Scekely and Rizzo, 2005); distance: Euclidean.

Potential spatial overlap between co-occurring mite taxa was investigated by calculating Pianka’s index and by using null-model approaches to suggest potential species interactions for more direct testing (Gotelli & Graves, 1996). We used the abundance of each taxon in the different body parts of RPW as a spatial resource. The niche plot of spatial resource utilisation showed the subelytral space as the region that contributed most to the high observed Pianka’s index, i.e. 0.854 (**Table 2; Fig. 5A**). The observed Pianka’s index was also significantly higher than the expected Pianka averages of 10000 simulated null-assemblages (*P* < 0.05), regardless of whether the RA2 (relaxed niche breadth) or RA3 (retaining niche breadth) random algorithm was used for their constructions **(Table 2**; **Fig. 5B**). The latter suggests that the observed distribution patterns of RPW-associated mite taxa were not randomly structured, but most likely occurred due to biological mechanisms that force the species to coexist.

**Table 2.** Estimates observed and expected values of Pianka’s niche overlap index. Expected values were determined using the RA2 and RA3 random algorithms to generate 10000 null assemblages. Significant *P*-value <0.05 is printed in bold.

| Index | Observed |  | Expected (RA2; 10000 replications) |  |  |  | Expected (RA3; 10000 replications) |  |  |  |
| --- | --- | --- | --- | --- | --- | --- | --- | --- | --- | --- |
|  | Estimate | Variance | Estimate | Variance | Lower tail | Upper tail | Estimate | Variance | Lower tail | Upper tail |
|  |  |  |  |  | *P*=(obs<exp) | *P*=(obs<exp) |  |  | *P*=(obs<exp) | *P*=(obs<exp) |
| Pianka | 0.854 | 0.013 | 0.768 | 0.002 | 0.975 | **0.024** | 0.343 | 0.004 | 0.999 | **<0.0001** |

**Figure 5.**
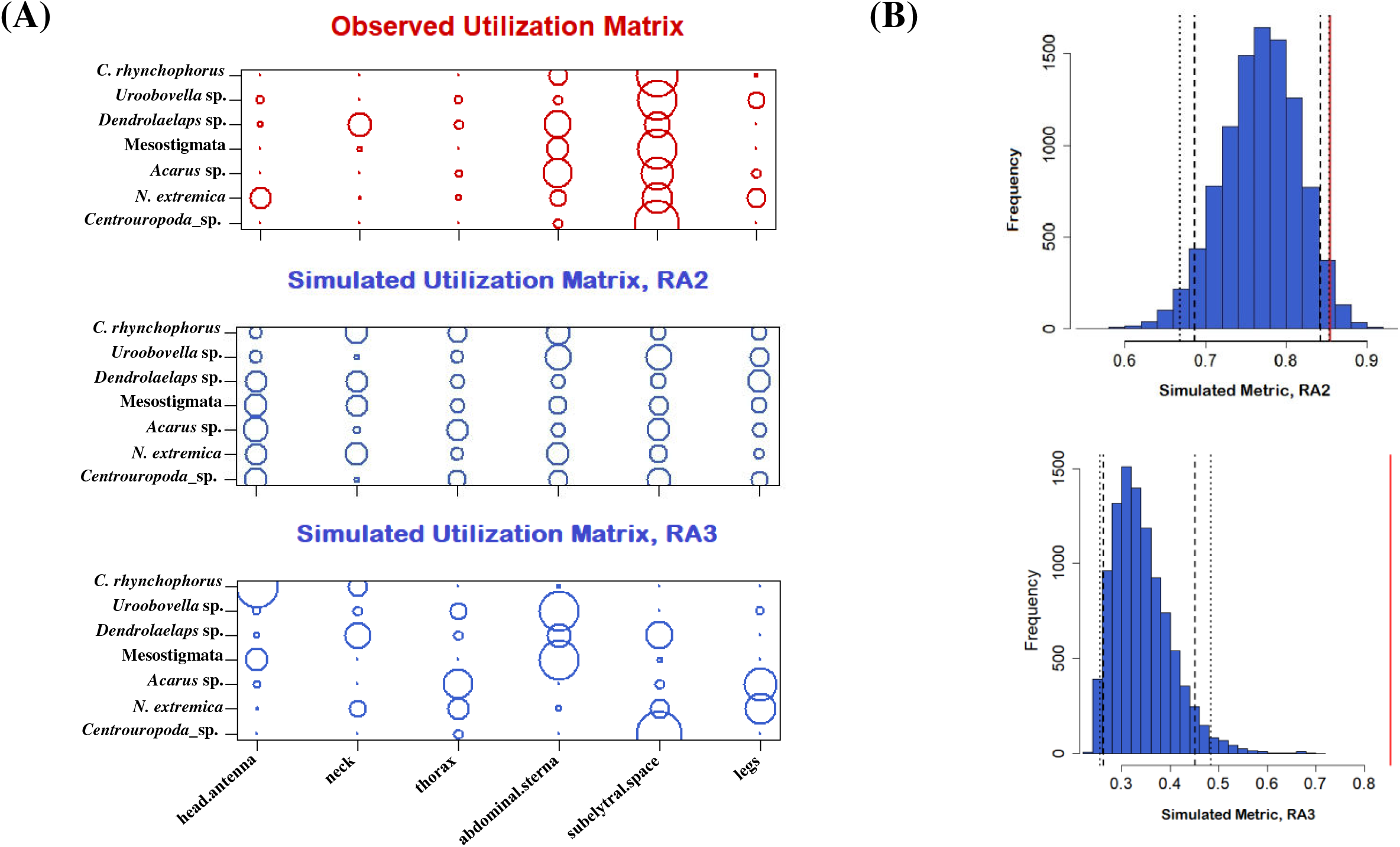
Null model analysis of phoretic niche overlap in red palm weevil-associated mite taxa implemented by the R package EcoSimR (Gotelli et al., 2015). (A) Niche plot and (B) histogram plot of resource utilisation (in our case red palm weevil body parts) by co-occurring phoretic mite taxa determined by the RA2 and RA3 random algorithms, each generating 10000 null assemblages. The niche plot is a visualisation of the mite taxa x body part utilisation matrix for the original data matrix (red) and the simulated (RA2 and RA3) data matrices (blue); the area of each circle is proportional to the utilisation of a body part by a mite taxon. The histogram shows the simulated Pianka values (blue bars) for each RA2 or RA3 algorithm, a vertical red line corresponding to the calculated Pianka metric for the original data, long dashed lines indicating the one-sided 95% confidence limits and short dashed lines indicating the two-sided 95% confidence limits.

### Interspecific coexistence of phoretic mites in the subelytral space increases as a function of body size difference with the dominant taxon

The interaction between pairs of taxa in the subelytral space was investigated using the Pearson correlation of their respective abundances in a network plot. Significant correlations were found between the taxa pairs *Centrouropoda* sp.-*Urobovella* sp. [r(236)=-0.13, *P*=0.046], *Centrouropoda* sp.-*C. rhynchphorus* [r(236)=0.19, *P*=0.004], *Centrouropoda* sp.-*N. extremica* [r(236)=-0.15, *P*=0.019], *Centrouropoda* sp.-*Acarus* sp. [r(236)=0.13, *P*=0.043], *Acarus* sp.- *Dendrolaelaps* sp. [r(236)=0.15, *P*=0.017] (**Fig. 6A**). Interestingly, the significant correlations between the dominant taxon *Centrouropoda* sp. and the common subordinate taxa *C. rhynchophorus* and *Acarus* sp., which were also the taxa with the smallest mites, were positive, whereas they were negative between *Centrouropoda* sp. and the sparse taxa *Urobovella* sp. And *N. extremica*, whose mites did not differ much in size compared to the dominant mites. A Pearson correlation coefficient was then calculated to assess the linear relationship between the difference in body size (Δ body size) and the difference in relative abundance in the subelytra (Δ relative abundance) for pairs of taxa formed between the dominant taxon *Centrouropoda* sp. and each of the other six taxa. The results showed a strong negative correlation between the two variables, r(4) =-0.85, *P*=0.033 (**Fig. 6B**).

**Figure 6.**
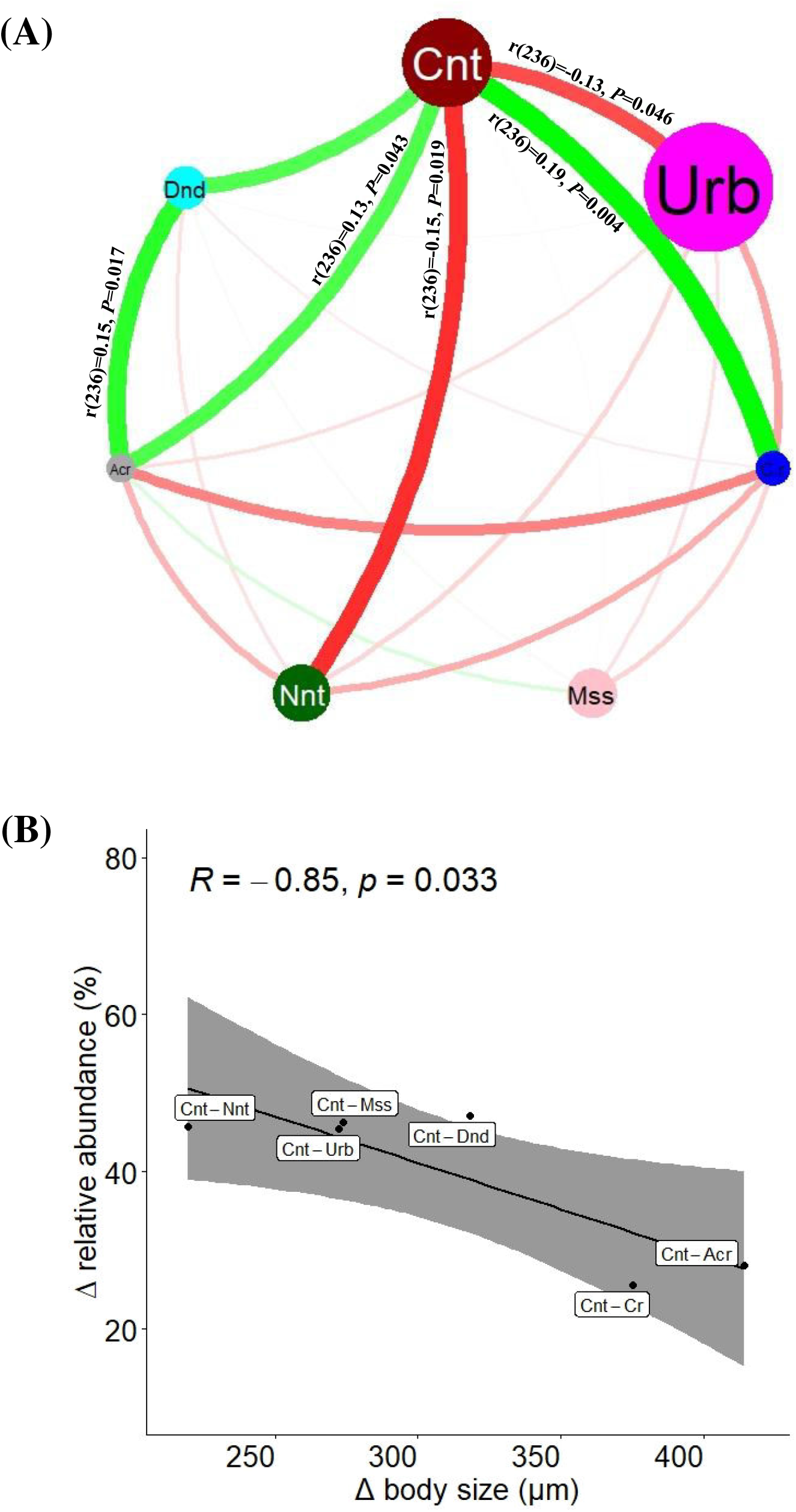
(A) Correlation network of mite taxa abundance in the subelytra of the 238 weevils examined. Each bubble node corresponds to one mite taxon. The size of the bubbles is proportional to the size of the mite taxon, i.e. Urb: *Uroobovella* sp. > Cnt: *Centrouropoda* sp. > Nnt: *Nenteria extremica* > Mss: Mesostigmata > Dnd: *Dendrolaelaps* sp. > C.r.: *Curculanoetus rhynchophorus* > Acr: *Acarus* sp. The green edges represent positive Pearson correlation coefficients (r > 0), while the red edges represent negative Pearson correlation coefficients (r < 0). The colour intensity and the width of the edges are proportional to r; the value of r is only shown for significant correlations (*P*<0.05). (B) Correlation between the difference in relative abundance (the percentage composition of mites of a given taxon to the total number of all taxa in the population) (Δ relative abundance) in the subelytral space and the difference in body size (Δ body size) of the dominant taxon (Cnt) with each of the other six mite taxa (i.e. Urb, Nnt, Mss, Dnd, C.r., Acr). The shaded area along the regression line corresponds to the 95% confidence intervals. The inset shows the values of r and *P*. Significance *P* <0.05.

## Discussion

Here we document a high species richness of phoretic deutonymphs associated with RPW in four districts of Northern Portugal —as part of a first study carried on these organisms in Portugal— and propose body size effects as determinants of coexistence or exclusion among mite taxa during the formation of distribution patterns on RPW. A total of seven mite taxa were found, but only two of them (i.e. *N. extremica* and *C. rhynchophorus*) could be identified to species level based on the original description of the life form of deutonymphs (Fain, 1974, Kontschán et al., 2014). In both cases, all described morphological details were found identical in the observed specimens. *N. extremica* has already been described in association with RPW (Dilipkumar et al., 2015). However, the first known association of *C. rhynchophorus* was with *Rhynchophorus phoenicis* (Fabricius, 1801) (Coleoptera: Curculionidae) (Fain, 1974) and later was also found associated with RPW in the United Arab Emirates (Al-Deeb et al., 2011). The other five taxa did not show sufficient similarity with the original descriptions and illustrations of the previously described species. Ongoing work in our laboratory attempts to classify and name the newly identified species.

The seven taxa were present in the four studied assemblages. However, their relative abundance (species evenness) varied between the districts, as did their diversity. The high number of mite taxa associated with RPW in Northern Portugal contrasts with the few species reported from other European countries where the invasive exotic RPW is also present, such as Italy (Longo & Ragusa, 2006; Mazza et al., 2011) and Spain (Abolafia and Ruiz-Cuenca, 2020), where only one taxon was reported, and Malta (Porcelli et al., 2009), where two taxa were reported. The reasons for this could be related to the different climatic conditions in the Euro-Mediterranean region and Northern Portugal, as mite communities have been shown to be very sensitive to temperature and precipitation fluctuations (Hassan et al., 2011; Kamczyc et al., 2022). In addition, some of the mite taxa identified in our work were very small (e.g. *C. rhynchophorus* and *Acarus* sp.) or occurred in low numbers (e.g. *N. extremica*, Mesostigmata, *Dendrolaelaps* sp. and *Uroobovella* sp.) and may have been overlooked in the other studies.

In our work, mites were found in all captured weevils and the prevalence of the seven taxa did not differ between districts or sex of the weevils, which indicates a high presence of these mite taxa in the Northern Portugal region and their close relationship with the RPW host. *Centrouropoda* sp. was the dominant taxon in the four studied assemblages. Phoretic species of this genus have shown strong adaptation to *Rhynchophorus* spp. in the past, e.g. *C. almerodai* to RPW in the Mediterranean countries of Italy, Malta, Tunisia and Egypt and the Southeast Asian countries of Malaysia and the Philippines (Dilipkumar et al., 2015) and *Centrouropoda* sp. to *R. Palmarum* in southern California (Gómez-marco et al., 2021). The other six taxa were less abundant and showed a high degree of intraspecific aggregation, especially the sparse taxa *Uroobovella* sp., *N. extremica*, Mesostigmata and *Dendrolaelaps* sp. This behaviour might be important to avoid exclusion by the dominant taxon, as coexistence of ecologically closely related species is possible if interspecific interference is not more important than intraspecific competition (den Boer, 1986; Ives, 1991). Nevertheless, aggregated distributions are also characteristic of parasites (Goater et al., 2014), and given the documented cases of transition from commensalism to antagonism in phoretic mites (Houck & Cohen, 1995; Seeman & Walter, 2023), there is a possibility that these taxa have a different relationship with RPW than phoresis, which needs further experimental verification.

We found the greatest amount of phoretic mites on the subelytral space of RPW. It has previously been suggested that this body region is preferred by phoretic mites because it can protect from fluctuating temperatures and ambient humidity but also to avoid being stripped as the weevil moves through dense palm fibres (Al-Deebet al., 2011; Dilipkumar et al., 2015). These authors speculated that the attachment of mites to body regions other than the subelytra might depend on the available space at the time of attachment. In our case, we observed mixed groups of the different taxa attaching to all body parts examined, especially in the subelytral space. Deciphering the stabilising mechanisms of the coexistence of mixed mite taxa on RPW could provide new insights into how species diversity is maintained in a multi-symbiont host. The use of null models helped us to determine a non-random structure of mite attachment pattern on RPW. For example, niche overlap among RPW-associated phoretic mites was significantly higher than expected by chance when the RA2 algorithm (relaxing niche breath) was used, suggesting that there are no restrictions on the utilisation of body parts, and also when the RA3 algorithm (retaining niche breath) was used, suggesting that some body parts may, however, be highly unsuitable, forcing taxa to coexist in other, more suitable parts, such as the subelytral space. As our study system consisted of different mite taxa within a considerable size range, we found it interesting to assess whether size regularity could mediate coexistence and exclusion mechanisms between taxa in the subelytral space. Our analysis was based solely on idiosoma length and did not consider the different morphologies of the seven taxa. The data presented here regarding the strong negative correlation between — body size and — relative abundance of the different taxa relative to the dominant *Centrouropoda* sp. in the subelytral space suggest that the distribution patterns of RPW-associated phoretic mites show body size-dependent effects that cause the dominant taxon to displace taxa of similar size and accept taxa with which it has the greatest size difference as co-habitants.

In summary, our results show a high diversity of phoretic deutonymphs associated with the invasive exotic pest RPW in northern Portugal, with seven taxa identified, of which *Centrouropoda* sp. was the most abundant and dominant taxon. Furthermore, the data presented here show that mite distribution patterns follow a non-random structure and that the interspecific coexistence of mixed mite taxa on RPW is significantly influenced by the difference in body size from the dominant taxon. These results may provide important insights into the use of phoretic mites in the biological control of RPW. Future work will investigate how the seasonal dynamics of individual taxa affect interactions between them and the host to gain further insight into the potential ecological and evolutionary consequences of their association with RPW.

## Data Accessibility

The datasets used and/or analysed during the current study are available from the corresponding author on reasonable request.

## Competing Interests Statement

The authors declare no conflict of interest. The funders had no role in the design of the study; in the collection, analyses, or interpretation of data; in the writing of the manuscript; or in the decision to publish the results.

## Author Contributions section

Conceptualization, C.A.-P., M.J.S., L.F.R., J.A.S.; methodology, M.J.S., C.A.-P., I.M., O.M.C.C.A., L.F.R., J.A.S.; investigation, I.M., D.S., J.O., C.G., R.A., N.P., F.C.; writing— review and editing, C.A.-P., M.J.S., I.M.; supervision, M.J.S., O.M.C.C.A., L.F.R., and C.A.-P.; project administration, C.A.-P., M.J.S.; funding acquisition, C.A.-P., M.J.S., L.R., J.A.S. All authors have read and agreed to the published version of the manuscript.

## Acknowledgements

This work has received financial support from the Portuguese national funds through FCT - Foundation for Science and Technology within the scope of PTDC/BIA-BMA/6363/2020, UIDB/04423/2020 442, UIDP/04423/2020, UIDP/50017/2020, UIDB/50017/2020, LA/P/0094/2020, by an FCT employment contract CEECIND/03501/2017 (to L.F.R.) and research fellowships PTDCASPPLA62282020-BI_LIC_2021_019 and 2022_083_BI_ASP-PLA (to I. M.). O.M.C.C.A. is funded by national funds (OE), through FCT, in the scope of the framework contract foreseen in the numbers 4, 5 and 6 of the article 23, of the Decree-Law 57/2016, of August 29, changed by Law 57/2017, of July 19.

## Notes

### Competing Interest Statement

The authors have declared no competing interest.

## References

Abolafia, J., & Ruiz-Cuenca, A. N. (2020). Phoretic invertebrates associated with Rhynchophorus ferrugineus (Coleoptera: Curculionidae) in Canarian date palm from southern Spain. Journal of Natural History, 54(35-36), 2265–2284. https://doi.org/10.1080/00222933.2020.1842930

Abo-Shnaf, R. I., & Allam, S. F. (2019). A new species of Centrouropoda (Acari: Uropodidae: Uropodina), with a key to the world species of the genus. Zootaxa, 4706(4), zootaxa. 4706.4704. 4701-zootaxa. 4706.4704. 4701.

Al-Deeb, M. A., Muzaffar, S. B., Abuagla, A. M., & Sharif, E. M. (2011). Distribution and abundance of phoretic mites (Astigmata, Mesostigmata) on Rhynchophorus ferrugineus (Coleoptera: Curculionidae). Florida Entomologist, 94(4), 748–755.

Al-Dhafar, Z. M., & Al-Qahtani, A. M. (2012). Mites Associated with the Red Palm Weevil, Rhynchophorus ferrugineus Oliver in Saudi Arabia with a Description of a New Species. Acarines, 6, 3–6.

Allam, S. F., & Elbadawy, A. R. (2017). Mass production of the facultative parasitic mite, Aegyptus rhynchophorus, as a natural enemy against the Red palm weevil in Egypt. VIII International Agriculture Symposium Jahorina, 5-8 October 2017 “AGROSYM 2017”. http://agrosym.ues.rs.ba/article/showpdf/BOOK_OF_PROCEEDINGS_2017_FINAL.pdf

Allam, S. F., Hassan, M. F., Taha, H. A., & Mahmoud, R. A. (2013). Hyperphoresy of phoretic Deutonymph of Aegyptus rhynchophorus (El-Bishlawi and Allam) (Acari: Uropodidae Trachyuropodidae) with the red palm weevil Rhynchophorus ferrugineus Olivier (Coleoptera: Curculionidae) in Egypt. Acarines, 7, 3–6

Allam, S. F. M., & El-Bishlawi, S. M. O. (2010). Description of Immature Stages of Aegyptus rhynchophorus (Elbishlawy & Allam), (Uropodina, Trachyuropodidae). Acarines, 4(1), 3–5. doi: 10.21608/ajesa.2010.163440

Bajerlein, D., Witaliński, W., & Adamski, Z. (2013). Morphological diversity of pedicels in phoretic deutonymphs of Uropodina mites (Acari: Mesostigmata). Arthropod Structure & Development, 42, 185–196.

Boavida, C., & Franca, M. F. (2008). Rhynchophorus ferrugineus (Olivier, 1790) (Coleoptera: Curculionoidea: Dryophthoridae), an exotic weevil recently introduced into Portugal. Boletín Sociedad Entomológica Aragonesa, 42, 425–426.

Cakmak, I., Hazur, S., Ulug, D., & Karagoz, M. (2013). Olfactory response of Sancassania polyphyllae (Acari: Acaridae) to its phoretic host larva killed by the entomopathogenic nematode, Steinernema glaseri (Rhabditida: Steinernematidae). Biological Control, 65, 212–217.

Castro, R., Santos, M. J. (2013). Metazoan ectoparasites of Atlantic mackerel, Scomber scombrus (Teleostei: Scombridae): macro- and microhabitat distribution. Parasitol Research, 112, 3579–3586.

Chao, A., Gotelli, N. J., Hsieh, T. C., Sander, E. L., Ma, K. H., Colwell, R. K., & Ellison, A. M. (2014). Rarefaction and extrapolation with Hill numbers: a framework for sampling and estimation in species diversity studies. Ecological Monographs, 84, 45–67.

Chao, A., Kubota, Y., Zelený, D., Chiu, C. -H., Li, C. -F., Kusumoto, B., Yasuhara, M., Thorn, S., Wei, C. -L., Costello, M. J., Colwell, R. K. (2020). Quantifying sample completeness and comparing diversities among assemblages. Ecological Research, 35(2), 292–314. https://doi.org/10.1111/1440-1703.12102

den Boer, P. J. (1986). The present status of the competitive exclusion principle. Trends in Ecology & Evolution, 1, 25–28. https://doi.org/10.1016/0169-5347(86)90064-9

DGAV. (2013). Plano de Ação para o controlo de Rhynchophorus ferrugineus (Olivier). Ministério da Agricultura e do Mar. Acessed at 15/06/2022. Retrieved from: https://www.drapc.gov.pt/base/documentos/plano_accao_r%20_ferrugineus%20_2013_dgav.pdf

Dilipkumar, M., Ahadiyat, A., Mašán, P., & Chuah, T. (2015). Mites (Acari) associated with Rhynchophorus ferrugineus (Coleoptera: Curculionidae) in Malaysia, with a revised list of the mites found on this weevil. Journal of Asia-Pacific Entomology, 18, 169–174.

Dormann, C. F. (2022). Using bipartite to describe and plot two-mode networks in R. https://cran.r-project.org/web/packages/bipartite/vignettes/Intro2bipartite.pdf

El-Sharabasy, H. M. (2010). A survey of mite species associated with the Red Palm Weevil, Rhynchophorus ferrugineus (Olivier) in Egypt. Egyptian Journal of Biological Pest Control, 20 (1), 67–70.

EPPO. (2023). [accessed 18 January 2023]. https://gd.eppo.int/taxon/RHYCFE.

Fain, A. (1974). Notes sur quelques hypopes d’Anoetidae. Bulletin et annales de la Société royale belge d’entomologie, 110, 58–68.

Faleiro, J. (2006) A review of the issues and management of the red palm weevil Rhynchophorus ferrugineus (Coleoptera: Rhynchophoridae) in coconut and date palm during the last one hundred years. International Journal of Tropical Insect Science, 26, 135–154.

Farahani, V. R. F., Ahadiyat, A., Mašán, P., & Dehvari, M. A. (2016). Phoretic uropodine mites (Acari: Mesostigmata) associated with the red palm weevil, Rhynchophorus ferrugineus (Coleoptera: Curculionidae) in Iran. Journal of Entomological and Acarological Research, 48, 5853.

Fernandes, D. (2016). Pragas no município do Porto: monitorização e proposta de gestão de três espécies de insetos. Master’s Thesis, Lisbon University. Retrieved from https://repositorio.ul.pt/bitstream/10451/25300/1/ulfc120675_tm_Diana_Fernandes.pdf

Fernandes, J. E. (2007). Memorando de 18 de Outubro 2007 sobre medidas da DRAP Algarve relativas a R. ferrugineus. Direcção-Regional de Agricultura e Pescas-Algarve.

Goater, T. M., Goater, C. P., & Esch, G. W. (2014). Parasitism: the diversity and ecology of animal parasites, 2nd edn. Cambridge, UK: Cambridge University Press.

Gobbin, T. P., Vanhove, M. P. M., Seehausen, O., & Maan, M. E. (2021). Microhabitat distributions and species interactions of ectoparasites on the gills of cichlid fish in Lake Victoria, Tanzania. International Journal for Parasitology, 51(2-3), 201–214. https://doi.org/10.1016/j.ijpara.2020.09.001

Gómez-Marco, F., Klompen, H., & Hoddle, M. S. (2021). Phoretic mite infestations associated with Rhynchophorus palmarum (Coleoptera: Curculionidae) in southern California. Systematic and Applied Acarology, 26, 1913–1926. https://doi.org/10.11158/saa.26.10.6

Gotelli, N. J., & Graves, G. R. (1996) Null models in ecology. Smithsonian Institution Press, Washington, 388 pp.

Gotelli, N. J., Hart, E. M., & Ellison, A. M. (2015) EcoSimR: Null model analysis for ecological data. R package. http://github.com/gotellilab/EcoSimR

Griffiths, D. A. (1970). A further systematic study of the genus Acarus L., 1758 (Acaridae, Acarina), with a key to species. Bulletin of the British Museum (Natur History Zoology), 85–118.

Hallett, R. H., Oehlschlager, A. C., & Borden, J. H. (1999). Pheromone trapping protocols for the Asian palm weevil, Rhynchophorus ferrugineus (Coleoptera: Curculionidae). International Journal of Pest Management, 45(3), 231–237.

Hassan, M., Nasr, A., Allam, S. F., Taha, H., & Mahmoud, R. A. (2011). Biodiversity and seasonal fluctuation of mite families associated with the red palm weevil, Rhynchophorus ferrugineus Oliver (Coleoptera: Curculionidae) in Egypt. Egyptian Journal of Biological Pest Control, 21(2), 317.

Helle, W., & Sabelis, M. W. (Eds.). 1985. Spider mites, their biology, natural enemies and control. Volume 1A. Elsevier. 405 pp.

Hodgkin, L., Elgar, M. A., & Symonds, M. R. E. (2010). Positive and negative effects of phoretic mites on the reproductive output of an invasive bark beetle. Australian Journal of Zoology, 58(3), 198–204.

Holt, R. D. (2009). Bringing the Hutchinsonian niche into the 21st century: Ecological and evolutionary perspectives. Proceedings of the National Academy of Sciences USA, 106, 19659– 19665. https://doi.org/10.1073/pnas.0905137106

Houck, M. A., & Cohen, A. C. (1995). The potential role of phoresy in the evolution of parasitism: radiolabelling (tritium) evidence from an astigmatid mite. Experimental And Applied Acarology, 19, 677–694.

Hsieh, T. C., Ma, K. H., & Chao, A. (2016) iNEXT: An R package for interpolation and extrapolation of species diversity (Hill numbers). Methods in Ecology and Evolution, 7, 1451.

Ives, A. R. (1991). Aggregation and coexistence in a carrion fly community. Ecological Monographs, 61, 75–94.

Jenő Kontschán, J., Mazza, G., Nannelli, R., & Roversi, P. F. (2014). Nenteria extremica n. sp., a new Uropodina mite (Acari Mesostigmata) collected on Rhynchophorus ferrugineus in Italy, with notes on other Uropodina mites associated with the Red Palm Weevil. Journal of Zoology, XCVII, 63–69.

Kamczyc, J., Dyderski, M. K., Horodecki, P., & Jagodziński, A. M. (2022). Temperature and precipitation affect seasonal changes in mite communities (Acari: Mesostigmata) in decomposing litter of broadleaved and coniferous temperate tree species. Annals of Forest Science, 79, 12. https://doi.org/10.1186/s13595-022-01129-9

Kinn, D. N. (1984). Life cycle of Dendrolaelaps neodisetus (Mesostigmata: Digamasellidae), a nematophagous mite associated with pine bark beetles (Coleoptera: Scolytidae). Environmental Entomology, 13(4), 1141–1144.

Krantz, G. W., & Walter, D. E. (2009). A Manual of Acarology (Third Edition ed.): Texas Tech University Press.

Lawlor, L. R. (1980) Structure and stability in natural and randomly constructed competitive communities. The American Naturalist, 116, 394–408. https://doi.org/10.1086/283634

Lindquist, E. E. (1975). Digamasellus Berlese, 1905, and Dendrolaelaps Halbert, 1915, with descriptions of new taxa of Digamasellidae (Acarina: Mesostigmata). The Canadian Entomologist, 107(1), 1–43.

Longo, S., & Ragusa, S. (2006). Presenza e diffusione in Italia dell’acaro Centrouropoda almerodai (Uroactiniinae Uropodina). Bollettino di Zoologia Agraria e di Bachicoltura (Ser. II), 38, 265–269.

Mazza, G., Cini, A., Cervo, R., & Longo, S. (2011). Just phoresy? Reduced lifespan in red palm weevils Rhynchophorus ferrugineus (Coleoptera: Curculionidae) infested by the mite Centrouropoda almerodai (Uroactiniinae: Uropodina). Italian Journal Of Zoology, 78(1), 101–105. https://doi.org/10.1080/11250003.2010.509135

McGraw, J. R., & Farrier, M. H. (1969). Mites of the superfamily Parasitoidea (Acarina-Mesostigmata) associated with Dendroctonus and Ips (Coleoptera: Scolytidae). Technical Bulletin. North Carolina Agricultural Experiment Station (192).

Mesbah H. A., Darwish E. T. E., Salem S. E., & Zayed T. M. (2008). Associations of three gamasid mite species with the red palm weevil, Rhynchophorus ferrugineus (Oliv.) in infested date palm farms in Beheira, Minufiya, Egypt. Journal of Agricultural Research, 33 (6), 1543–1551.

Morrill, A., Nielsen, Ó. K., Skírnisson, K., & Forbes, M. R. (2022). Identifying sources of variation in parasite aggregation. Peer J, 10, e13763. http://doi.org/10.7717/peerj.13763

Murphy, S. T., & Briscoe, B. R. (1999). The red palm weevil as an alien invasive: biology and the prospects for biological control as a component of IPM. Biocontrol News and Information, 20, 35–46.

Patil, G. P. & Taillie, C. (1979) An Overview of Diversity. In: Grassle, F., Patil, G.P., Smith, W. & Taillie, C., Eds., Ecological Diversity in Theory and Practice. International Co-Operative Publishing House, Fairland, 3–27.

Pete, N. (2010). Rhynchophorus ferrugineus in Europe survey results. Internacional Conference on Red Pal Weevil (p. 37). Valencia - Spain: UE.

Pianka, E. R. (1974). Niche overlap and diffuse competition. Proceedings of the National Academy of Sciences USA, 71, 2141–2145.

Porcelli, F., Ragusa, E., D’Onghia, A. M., Mizzi, S., & Mifsud, D. (2009). Occurrence of Centrouropoda almerodai and Uroobovella marginata (Acari: Uropodina) phoretic on the Red Palm Weevil in Malta. Bulletin of the Entomological Society of Malta, 2, 61–66.

Poulin, R. (1993). The disparity between observed and uniform distributions: a new look at parasite aggregation. International Journal for Parasitology, 23, 931–944. https://doi.org/10.1016/0020-7519(93)90060-C

Qin, Z., Li, D., Luo, Y., Huang, X., Goebel, F. R., & Zhou, Z. (2022). First record of damage by the red palm weevil, Rhynchophorus ferrugineus (Olivier, 1790) in sugarcane fields in China. International Journal of Pest Management, DOI: 10.1080/09670874.2022.2145521.

R Core Team. R: A Language and Environment for Statistical Computing. R Foundation for Statistical Computing, Vienna, Austria. 2021. Available online: http://www.R-project.org/ (accessed on 19 September 2021).

Ramos, A. P., Caetano, M. F., Rocha, M., & Lima, S. B. (2013). Doenças e pragas que condicionam o uso de palmeiras em espaços verdes. Revista da Associação Portuguesa de Horticultura, 112, 37–40.

Rohde, K. (1984). Ecology of marine parasites. Helgolander Meeresunters, 37, 5–33. https://doi.org/10.1007/BF01989293

Rózsa, L., Reiczigel, J., & Majoros, G. (2000). Quantifying parasites in samples of hosts. Journal of Parasitology, 86, 228–232.

Seeman, O. D., & Walter, D. E. (2023). Phoresy and mites: more than just a free ride. Annual Review of Entomology, 68, https://doi.org/10.1146/annurev-ento-120220-013329

Slimane-Kharrat, S., & Ouali, O. (2019). Mites associated with the red palm weevil (Rhynchophorus ferrugineus) in Tunisia. Tunisian Journal of Plant Protection, 14 (2), 29–38.

Stirling, G. R., Stirling, A. M., & Walter, D. E. (2017). The Mesostigmatid mite Protogamasellus mica, an effective predator of free-living and plant-parasitic nematodes. Journal of Nematology, 49(3), 327–333.

Suzuki, R., & Shimodaira, H. (2006). Pvclust: an R package for assessing the uncertainty in hierarchical clustering. Bioinformatics, 22(12), 1540–1542. https://doi.org/10.1093/bioinformatics/btl117

Wisniewski, J., Hirschmann, W., & Hiramatsu, N. (1992). New species of Centrouropoda (Uroactiniinae, Uropodina) from Philippines, Brazil and Middle Africa. Acarologia, 33(4), 313–320.

Womersley, H. (1954). Two new species of mites (Acarina: Mesostigmata: Ascidae) associated with bark-boring beetles from South Australia. Records of the South Australian Museum, 11, 113–116.

Zeileis, A., & Kleiber, C. (2014). Ineq: Measuring Inequality, Concentration, and Poverty. R package. https://cran.r-project.org/web/packages/ineq/ineq.pdf

